# High temperatures augment inhibition of parasites by a honey bee gut symbiont

**DOI:** 10.1101/2023.03.20.533504

**Authors:** Evan C Palmer-Young, Lindsey M Markowitz, Wei-Fone Huang, Jay D Evans

## Abstract

Temperature affects growth, metabolism, and interspecific interactions in microbial communities. Within animal hosts, gut bacterial symbionts can provide resistance to parasitic infections. Infection can also be shaped by host body temperature. However, the effects of temperature on the antiparasitic activities of gut symbionts have seldom been explored. The *Lactobacillus*-rich gut microbiota of facultatively endothermic honey bees is subject to seasonal and ontogenetic changes in host temperature that could alter the effects of symbionts against parasites. We used cell cultures of a *Lactobacillus* symbiont and an important trypanosomatid gut parasite of honey bees to test the potential for temperature to shape parasite-symbiont interactions.

We found that symbionts showed greater heat tolerance than parasites and chemically inhibited parasite growth via production of acids. Acceleration of symbiont growth and acid production at high temperatures resulted in progressively stronger antiparasitic effects across a temperature range typical of bee colonies. Consequently, the presence of symbionts reduced both peak growth rate and heat tolerance of parasites. Results suggest that the endothermic behavior of honey bees could potentiate the effectiveness of gut symbionts that limit parasites’ ability to withstand high temperature, implicating thermoregulation as a reinforcer of core symbioses and possibly microbiome-mediated antiparasitic defense.

**IMPORTANCE:** Two factors that shape the resistance of animals to infection are body temperature and gut microbiota. However, temperature can also alter interactions among microbes, raising the question of whether and how temperature changes the antiparasitic effects of gut microbiota. Honey bees are agriculturally important hosts of diverse parasites and infection-mitigating gut microbes. They can also socially regulate their body temperatures to an extent unusual for an insect. We show that high temperatures found in honey bee colonies augment the ability of a gut bacterial symbiont to inhibit growth of a common bee parasite and reduce the parasite’s ability to grow at high temperatures. This suggests that fluctuations in colony and body temperatures across life stages and seasons could alter the protective value of bees’ gut microbiota against parasites, and that temperature-driven changes in gut microbiota could be an underappreciated mechanism by which temperature— including endothermy and fever— alters animal infection.

## INTRODUCTION

Temperature has a fundamental effect on biological rates that influence organismal physiology and interspecific interactions (1–4), including infection of hosts by parasites (5–8). The influence of temperature on infection outcome reflects not merely the direct effects of temperature on the performance of parasites, but on parasites’ performance relative to that of the host and its immune system (6, 8). In many hosts, attainment of high internal temperatures can reduce the intensity and effects of infection (9–13), whether due to direct inactivation of parasites or potentiation of host immunity (10, 14–16). Such temperature-mediated increases in host resistance to infectious disease could favor the evolution of energetically expensive, endothermic life history strategies characterized by sustained high body temperatures (17).

In addition to the defenses of the host itself, parasites that establish in the gut are also influenced by cooccurring species that form the host gut microbiota and are likewise influenced by host temperature. The community of non-pathogenic microbial species (hereafter referred to as ‘symbionts’, although this term *sensu stricto* includes all host-associated species) in the host gut can substantially influence infection outcome via physical, chemical, and host immune system-mediated effects (18). A growing number of studies have indicated that temperature influences the composition of the gut symbiont community in ways that are qualitatively similar across distantly related hosts (19). However, despite the known effects of temperature on microbial community composition, biochemistry, and the microbiome and parasitic infection of hosts, the ways in which temperature alters the effects of gut symbionts on parasites has received little attention (19). A more complete knowledge of this area would elucidate how climate and host thermoregulation, including the constitutively high body temperatures of endotherms and infection-associated elevations in body temperature known as ‘fever’, could affect parasitism via symbiont-driven effects.

Honey bees and their associated microbiota offer an excellent opportunity to study the joint effects of temperature and gut symbionts on parasites. Honey bees are facultatively endothermic, capable of sustaining body and hive temperatures that are similar to those of mammals. Like other insects, however, they are subject to wide temperature variations during outdoor foraging and in the winter (20, 21). Honey bees also possess a broadly consistent, culturable gut bacterial community (22, 23) that can improve host resistance to parasites (24) that can threaten colony health (25–27).

Trypanosomatid parasites of the honey bee hindgut, namely *Crithidia mellificae* (28) and the currently dominant *Lotmaria passim* (29), are prevalent, widespread, and in some cases associated with shortened bee lifespan and colony losses (26, 30–36). These honey bee-associated species are close relatives of the bumble bee parasite *Crithidia bombi. Crithidia bombi* infection is profoundly affected by the bumble bee gut microbiota (37, 38), which is taxonomically similar to that of honey bees (39). This includes negative correlations with the abundance of *Lactobacillus* ‘Firm-5’ clade members (38, 40), which colonize the hindgut of honey bees as well (39, 41–43). Bumble bee-associated *Lactobacillus* symbionts exerted temperature-dependent, acidity-driven effects on *C. bombi* growth in cell cultures (44, 45), and could contribute to the reduction of infection seen in bees reared at high temperatures (37 °C (46)). The gut microbiota of other insects can also influence infection with trypanosomatids, including species of medical and veterinary importance (47). However, the effects of the honey bee microbiome, including Lactobacilli, on trypanosomatid infection remain unclear. The only work thus far that tested the effects of gut microbial manipulations on infection found that pre-treatment of worker bees with another symbiont, *Snodgrassella alvi,* resulted in greater intensities of *L. passim* infection (48).

To evaluate how temperature affects growth rates of honey bee *Lactobacillus* symbionts relative to trypanosomatid parasites and the effects of symbionts on parasite growth, we conducted three sets of experiments to: (1) Compare the temperature dependence of parasite and symbiont growth rates; (2) test for effects of symbionts and the acids they produce on parasite growth; and (3) determine how the effects of symbionts vary with temperature and alter the thermal niche of parasites. Our results show that the high colony temperatures produced by the endothermic behavior of bees can selectively favor the growth of symbionts and their inhibition of parasites, such that the presence of symbionts effectively reduces parasite heat tolerance.

## MATERIALS AND METHODS

### Cell Cultures

Two strains of *Lactobacillus* nr. *melliventris* (49), kindly provided by N.J. Moran and J.E. Powell, were grown in De Man, Rogosa and Sharpe (MRS) media supplemented with 0.05% cysteine in screw-cap microcentrifuge tubes at 37 °C. These isolates are hereafter referred to by full genus and strain names, for consistency with their original description and to differentiate their genus from *Lotmaria.* The honey bee trypanosomatid parasites *C. mellificae* (ATCC 30254 (28)) and *L. passim* (strain BRL (29)) were obtained from the American Type Culture Collection and R.S. Schwarz. Trypanosomatids were grown in “FPFB” medium including 10% heat-inactivated fetal bovine serum (pH 5.9-6.0 (50)) at 28 °C in vented cell culture flasks. Cultures were transferred to fresh media every 2 d.

### Effects of temperature on symbiont vs. parasite growth rate

Dense cultures (net OD ∼0.8) were diluted to a starting OD of 0.010 in fresh media and incubated in microcentrifuge tubes at 3 °C increments between 25 and 46 °C. We ran six experimental blocks, each of which used parallel incubations at 7 different temperatures (3 blocks in 3 °C increments from 25-43 °C and 3 blocks from 28-46 °C). One replicate tube per strain and temperature was destructively sampled at each of two time points (18 and 24 h). Growth rates were calculated as the slope of the curve of ln(OD) vs. time using the 18 h time point only, at which point we could be confident that cultures were still in the logarithmic phase of growth. For comparison, we used previously reported growth rates of the two honey bee trypanosomatids (51), which included two replicate runs per temperature at 20 °C and in 2 °C increments from 23-41 °C, plus a third replicate for each of four temperatures (25, 31, 33, and 35 °C).

We modeled the temperature dependence of growth rate for each *Lactobacillus* and trypanosomatid strain using a Sharpe-Schoolfield equation modified for high temperatures (15, 52, 53).

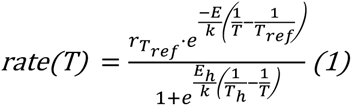

In Equation (1), *rate* refers to the maximum specific growth rate (in [h^-1^]); *r_Tref_* is the growth rate (in [h^-1^]) at the calibration temperature *T_ref_* (293K, i.e., 20°C, chosen to be well below the temperature of peak rate)); *E* is the activation energy (in eV), which primarily affects the upward-sloping portion of the thermal performance curve (i.e., responsiveness of growth to temperature) at suboptimal temperatures; *k* is Boltzmann’s constant (8.62·10^-5^ eV·K^−1^); *E_h_* is the deactivation energy (in eV), which determines how rapidly the thermal performance curve decreases at temperatures above the temperature of peak growth *T_pk_* (in K); *T_h_* is the high temperature (in K) at which growth rate is reduced by 50% (relative to the value predicted by the Arrhenius equation—which assumes a monotonic, temperature-dependent increase) (53); and *T* is the experimental incubation temperature (in K). Four models were fit, one for each strain of *Lactobacillus* and each trypanosomatid (51).

Models were optimized using nonlinear least squares, implemented with R packages rTPC and nls.multstart (54). Confidence intervals on parameter values and predicted growth rates were obtained by bootstrap resampling of the residuals (10,000 model iterations, R package “car” (55)). Expected ratios between growth rates of symbionts and parasites were calculated by dividing the average of the predicted growth rates for the two *Lactobacillus* strains by the predicted rates for each of the trypanosomatid species at the corresponding temperature. We also used the bootstrap model predictions to estimate the following traits not explicitly included in the model: peak growth rate; temperatures of peak growth rate (*T_pk_*), and 50% inhibition relative to the peak value due to low and high temperatures (referred to as “cold tolerance” and “heat tolerance” in figures); and thermal niche breadth (i.e., the number of degrees between the low and high temperatures of 50% rate inhibition). The 0.025 and 0.975 quantiles for parameter estimates, predicted growth rates at each temperature, and traits derived from bootstrap predictions were used to define 95% confidence intervals. Estimates from different models were considered significantly different from each other when their 95% confidence intervals did not overlap. This and all subsequent analyses were conducted using R for Windows v4.0.3 (56).

### Effects of symbiont spent medium and acidity on parasite growth

To generate spent media, each of the two *Lactobacillus* strains was grown in 15 mL conical centrifuge tubes for 2 d at 37 °C from a 100x dilution of stationary phase cultures. The resulting supernatant (pH 4.6) was sterile-filtered, and an aliquot was neutralized with 1M NaOH to the original pH of the MRS media (pH 5.6). Meanwhile, an aliquot of fresh MRS media was acidified with glacial acetic acid (the main acid in the honey bee gut (57) and of isolated *Lactobacillus melliventris* under aerobic conditions (57)) to the pH of the spent media (pH 4.6). Each of the two trypanosomatids was then grown in a mixed growth medium consisting of equal volumes of fresh trypanosomatid-specific “FPFB” media and one of six *Lactobacillus-*specific MRS media-based treatments: spent or neutralized media from each of the two *Lactobacillus* strains, acidified MRS media, or a fresh MRS media control. The spent media treatments tested for symbiont-mediated inhibition of parasite growth; the acidified fresh media and neutralized spent media treatments indicated the extent to which this inhibition was dependent on pH.

To assess parasite growth, cell cultures of each trypanosomatid were diluted to a net OD of 0.040 in the appropriate mixed media. Growth of n = 6 replicate wells per treatment was measured in 96-well plates incubated in a plate reader spectrophotometer at 31 °C. We included cell-free negative controls for each of the MRS media treatments, to control for changes in OD of media independent of parasite growth. OD values were corrected by subtraction of the OD of the corresponding cell-free media treatment and time point. Growth was estimated by measurement of optical density (OD) at 600 nm in 5 min intervals for 24 h, with a 30 s shake before each read. Growth rate was calculated as the maximum slope of ln(OD) vs. time over a rolling 4 h window, after exclusion of the first 3 h to allow OD to stabilize (15, 51, 58). Samples for which the r-squared value for the growth rate fell below 0.9 (vs. >0.98 for all non-acidified samples) were considered to have a negligible growth rate and were assigned a rate of 0. Visual inspection of growth curves indicated that none of these samples exceeded a net OD of 0.050.

### Temperature-dependent effects of symbionts on parasite growth in parasite-symbiont cocultures

To evaluate how temperature modulates the ability of symbionts to inhibit parasites, we grew *L. passim* in the presence and absence of *Lactobacillus* strain wkB10 at temperatures between 20.7 and 38 °C, which spans the range of temperatures commonly observed in worker bees within the winter cluster of colonies in a temperate climate (21). A dense culture of *L. passim*, grown 2 d at 28 °C from a 25-fold dilution, was diluted to a net OD of 0.020 in fresh FPFB media. A dense culture of *Lactobacillus* strain wkB10, grown 2 d at 37 °C from a 100-fold dilution, was diluted to a net OD of 0.040 in fresh MRS media. The *L. passim* cell suspension was then mixed with an equal volume of either the *Lactobacillus* cell suspension (“*Lactobacillus* present” treatment, initial net OD of 0.010 for the parasite and 0.020 for the symbiont) or fresh MRS without cells (“*Lactobacillus* absent”). These initial densities were chosen such that *L. passim* would remain in log-phase growth throughout the experiment, while *Lactobacillus* would approach stationary phase at the higher temperatures.

Two replicate microcentrifuge tubes, each containing 1400 μL of media, were incubated in parallel for 23 h at each of 7 different temperatures; two additional tubes were refrigerated at ∼0 °C immediately following setup for measurement of initial cell densities and media pH. Following the incubation, parasite densities were quantified by microscopic cell counts using a Neubauer hemocytometer at 400x magnification, using a 100 μL aliquot of each tube diluted with an equal volume of 50% glycerol to slow the movement of the parasite cells. Parasite growth rates were estimated based on the ratio of parasite cell densities at 23 h vs 0 h of incubation. The remaining sample volume was used for measurement of pH. The experiment was repeated 3 times for a total of 6 replicate samples per temperature and *Lactobacillus* treatment.

To quantify the temperature dependence of parasite growth rate, separate Sharpe-Schoolfield models were fit to the data for parasite growth rate in the presence and absence of *Lactobacillus*, using the approach described above (<u>Methods: Effects of temperature on symbiont vs. parasite growth rate</u>). To compute the proportion of parasite growth inhibition by *Lactobacillus*, we averaged growth rates of the two *Lactobacillus*-containing and *Lactobacillus*-free replicates within each temperature and experimental block. Percent inhibition was then computed from the ratios of growth rates in the presence vs. absence of *Lactobacillus* for each temperature-block combination.

We modeled the correlations between temperature and parasite inhibition, temperature and pH, and pH and parasite inhibition using generalized linear models (59). Each model included experiment block as a random effect. In addition, the models for the effects of temperature and of pH on parasite inhibition included quadratic predictor terms (i.e., (temperature)^2^, (pH)^2^) to better describe the curvilinear relationship between these variables that was apparent from visual inspection of the data. Models were fit using R package glmmTMB (59), with predictions generated with R package emmeans (60). Graphs were produced using R packages ggplot2 and cowplot (61, 62).

## RESULTS

### Effects of temperature on growth rates

Relative to the trypanosomatid parasites measured in our previous experiments (51)), growth of both strains of *Lactobacillus* symbionts increased more strongly with temperature and peaked at higher temperatures, resulting in progressively higher projected growth rates of the symbionts relative to the parasites--particularly *L. passim—*as temperatures increased **(Figure 1**). *Lactobacillus* growth rates increased by 6- to 7-fold over the 15 °C temperature window between 25 and 40 °C, whereas growth rates of trypanosomatids varied by <2-fold over a similar 15 °C interval from 20 to 35 °C. These differences in temperature responsiveness were reflected in the estimates for activation energy (model parameter *E*), which was more than twice as high in both strains of *Lactobacillus t*han in either species of trypanosomatid (**Figure 2;** see **Supplementary Figure 1** for additional parameters). In addition, *Lactobacillus g*rowth rates peaked at temperatures more than 4 °C higher than those of *C. mellificae* and more than 6 °C higher than those of *L. passim* **(Figure 2)**. The more steeply angled, high temperature-shifted thermal performance curves of the two Lactobacilli were likewise reflected by their higher temperatures of cold- and heat-related 50% growth inhibition and narrower thermal breadth (i.e., width of the thermal interval over which growth rates exceeded 50% of the peak value**, Supplementary Figure 2**).

**Figure 1.**
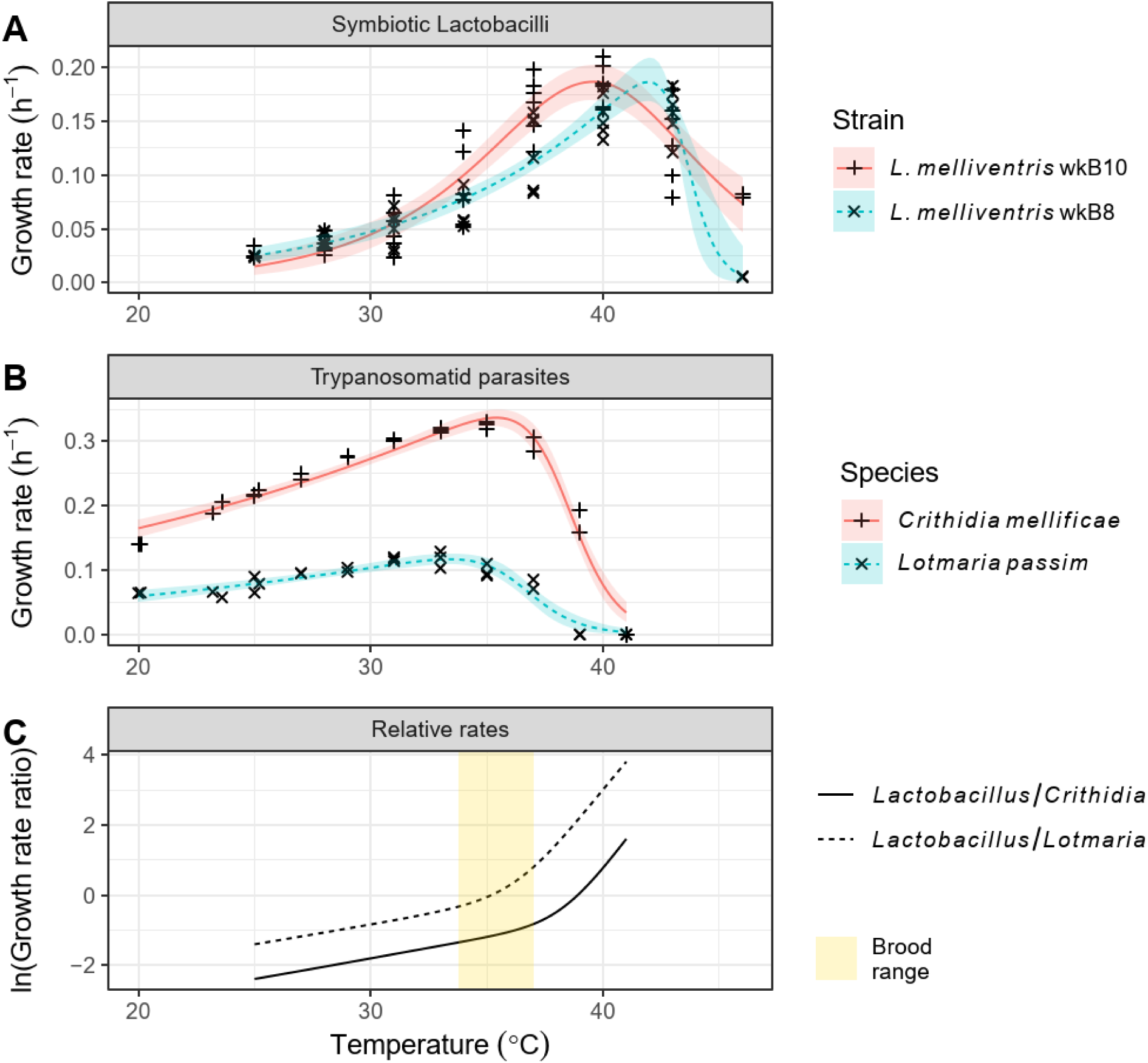
Growth of *Lactobacillus melliventris* honey bee gut symbionts responds more strongly to temperature and persists at higher temperatures than growth of the trypanosomatid parasites *Crithidia mellificae* and *Lotmaria passim*. **(A)** Growth rates of *Lactobacillus* strains wkB8 (solid lines) and wkB10 (dashed lines) between 25 and 46 °C. Points show observed rates. Lines and shaded bands show Sharpe-Schoolfield model predictions and 95% bootstrap confidence intervals. **(B)** Sharpe-Schoolfield models for temperature-dependent growth of *C. mellificae* (solid lines) and *L. passim* (dashed lines), as reported previously (51). **(C)** Projected growth rates of *Lactobacillus* (average of strains wkB8 and wkB10) relative to *C. mellificae* (solid lines) and *L. passim* (dashed lines) between 25 and 41 °C. Yellow shaded region represents temperature range at center of brood-rearing honey bee colony (33.8-37 °C (20)).

**Figure 2.**
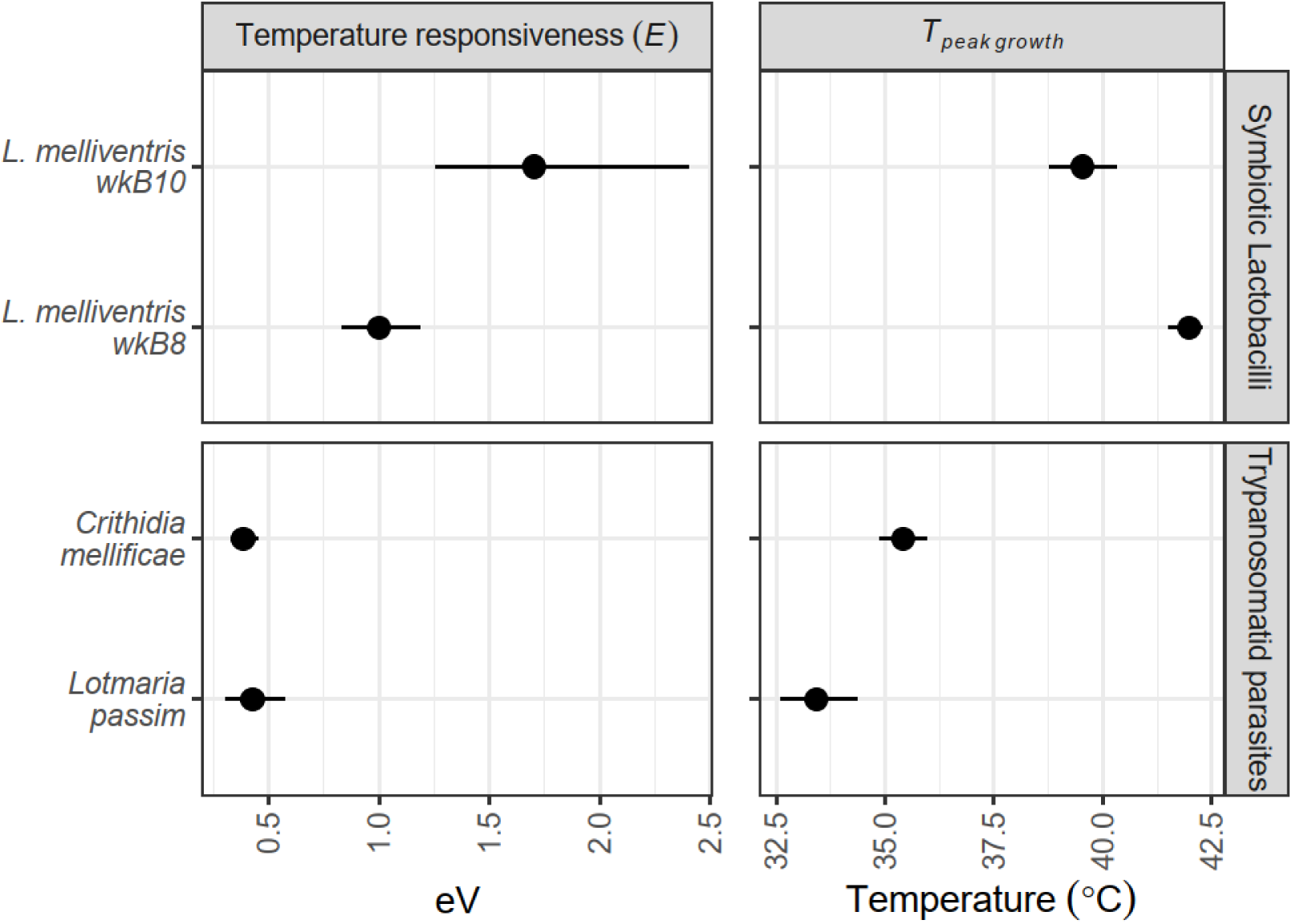
Growth rates of symbiotic Lactobacilli (upper panels) were more responsive to temperature and peaked at higher temperatures than did growth of trypanosomatid gut parasites (lower panels). Left panels show Sharpe-Schoolfield model parameter *E*, a measure of how strongly growth rate increases with temperature at temperatures below the temperature of peak growth (right panels). Points and error bars show bootstrap medians and 95% confidence intervals.

Based on model predictions, the expected growth rates of symbionts relative to trypanosomatids increased with temperature across the 25-37 °C interval that is most common in honey bee workers (21), changing by 4.67-fold for *Lactobacillus* (averaged across the two strains) relative to *C. mellificae* and 9.25-fold relative to *L. passim* **(Figure 1**). Even over the narrow 33.8-37 °C temperature range of the central honey bee brood cluster (20), growth rates of *Lactobacillus* increased by 65% relative to *C. mellificae* and by more than 3-fold relative to *L. passim* (**Figure 1**).

### Acidity-mediated inhibitory effects of *Lactobacillus* spent media

Spent media from each of the two *Lactobacillus* strains resulted in full inhibition of both *C. mellificae* and *L. passim* **(Figure 3)**. Acidification of fresh media to an equivalent pH with acetic acid likewise extinguished growth, whereas neutralization of the spent media restored 89-90% of growth in *C. mellificae* and 81-82% in *L. passim,* indicating that acidity was necessary and sufficient to account for most of the observed inhibition by spent media **(Figure 3)**.

**Figure 3.**
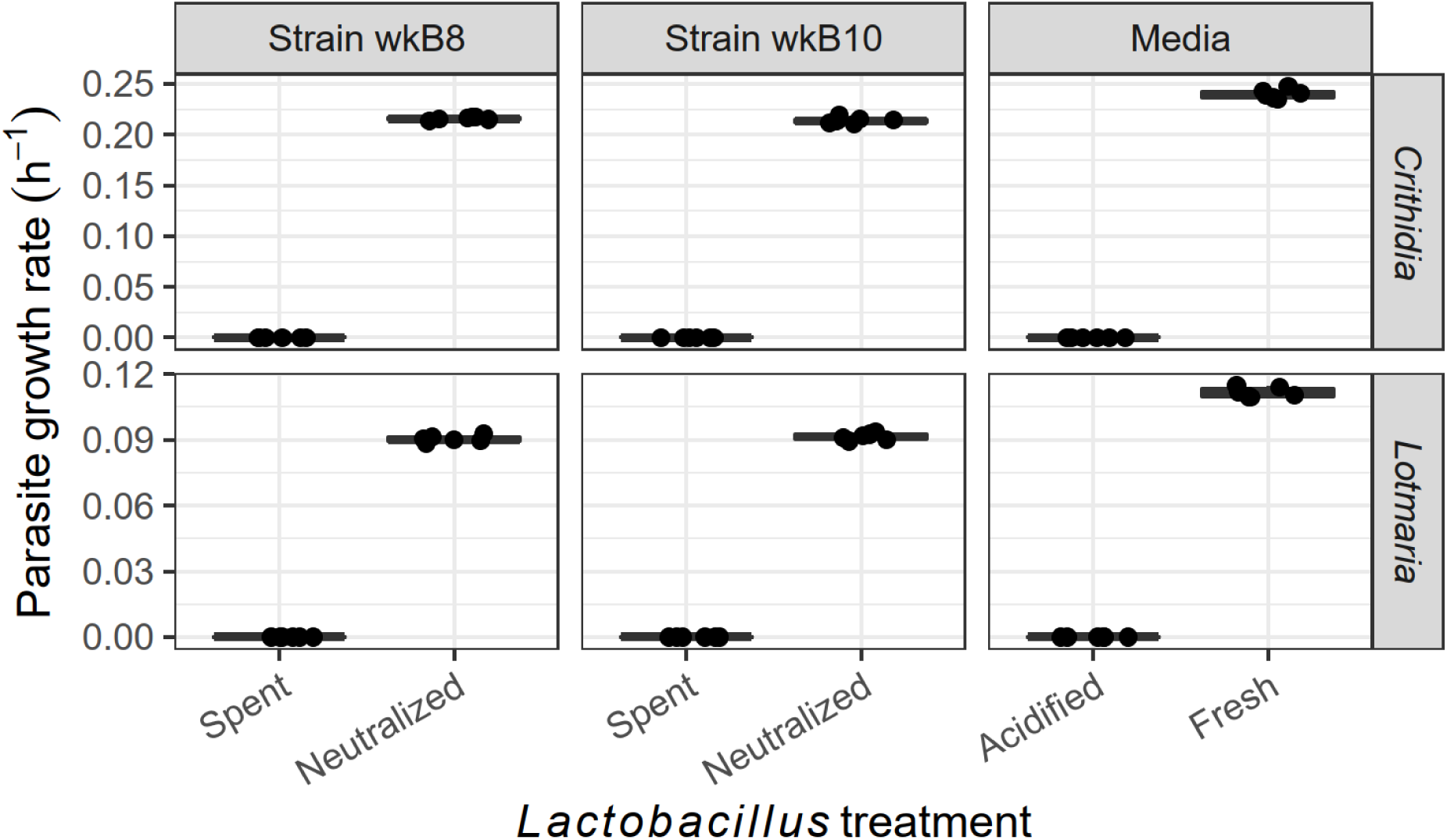
Acidity-mediated inhibition of *Crithidia mellificae* and *Lotmaria passim* growth by *Lactobacillus* spent media. Growth rates of *C. mellificae* (upper panels) and *L. passim* (lower panels) in spent media (A, C) of *Lactobacillus melliventris* strains wkB8 (left) and wkB10 (center) before and after neutralization from pH 4.6 to 5.6 with 1M NaOH, and in fresh *Lactobacillus* media (right) with and without acidification from pH 5.6 to 4.6 with acetic acid. Boxplots show medians and interquartile range for n = 6 samples per group. Points show individual observations, horizontally offset to mitigate overplotting.

### Temperature-dependent effects of *Lactobacillus* on *Lotmaria passim* growth in parasite-symbiont cocultures

In parasite-symbiont cocultures, the presence of *Lactobacillus* altered the thermal niche of *L. passim* by inhibiting the parasite’s growth in a temperature-dependent fashion **(Figure 4**). The 95% confidence intervals for model-predicted parasite growth rates in the presence and absence of *Lactobacillus* overlapped below 23 °C, but the symbiont’s presence had increasingly pronounced inhibitory effects as temperature increased **(Figure 4**). These effects resulted in a 24% lower predicted peak parasite growth rate and a 1.9 °C reduction in the temperature at which parasite growth declined to less than 50% of peak rate, from 34.68 °C in the absence of *Lactobacillus* (95% CI: 34.16-35.55 °C) to 32.76 °C in its presence (95% CI: 32.06-33.46 °C) **(Figure 5)**. Confidence intervals for other parameters and traits overlapped in the two *Lactobacillus* treatments **(Supplementary Figures 3 and 4**).

**Figure 4.**
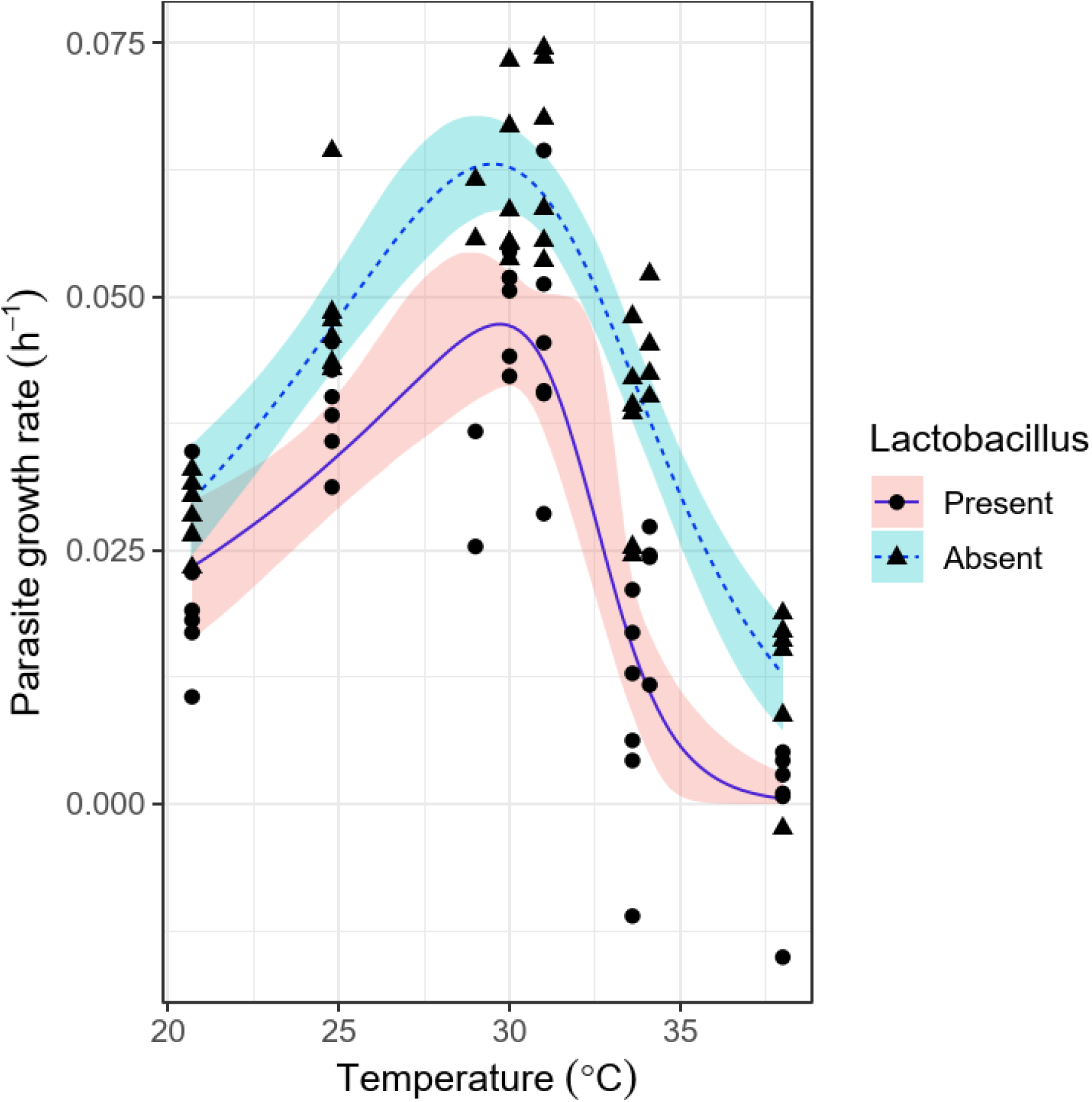
Coculture with *Lactobacillus* affects *Lotmaria passim* growth in a temperature-dependent manner, with stronger effects of the symbiont at higher temperatures. Lines and shaded bands show Sharpe-Schoolfield model fits and 95% confidence intervals for *L. passim* growth rate in the presence (solid lines) and absence (dashed lines) of *Lactobacillus*. Points show individual observations; circles and triangles indicate presence and absence of *Lactobacillus,* respectively.

**Figure 5.**
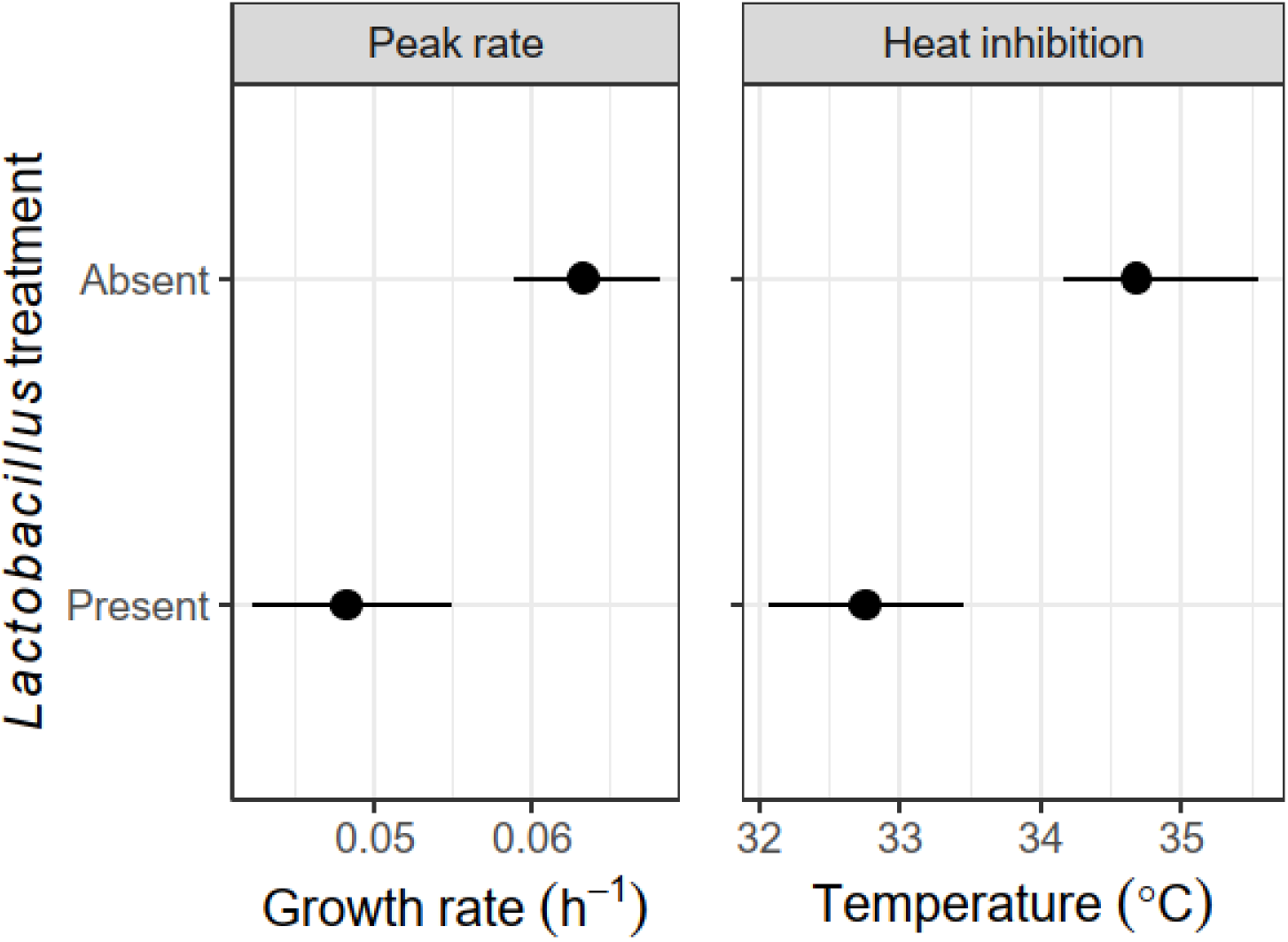
The presence of *Lactobacillus* reduced peak *Lotmaria passim* growth rate (left) and lowered the temperature at which growth rate declined to <50% of peak value (right). Points and error bars show estimates and 95% bootstrap confidence intervals for traits derived from Sharpe-Schoolfield model fits shown in Figure 4.

The inhibitory effects of *Lactobacillus* increased quadratically with temperature (Temperature: coefficient -25.71 ± 8.82 SE, Z = -2.91, P = 0.004; Temperature^2^: coefficient 0.51 ± 0.15 SE, Z = 3.39, P < 0.001); inhibition exceeded 50% at 32.9 °C (95% CI: 30.26-34.82 °C, **Figure 6A**). The final pH of the coculture decreased by 0.098 ± 0.0060 SE pH units per 1 °C increase in temperature (Z = -16.38, P < 0.001, **Figure 6B**). Across all temperatures, stronger inhibitory effects of *Lactobacillus* were quadratically correlated with lower final pH (pH: coefficient -707.75 ± 203.32 SE, Z = -3.48, P < 0.001; pH^2^: coefficient 65.60 ± 20.05 SE, Z = 3.27, P = 0.001), with 50% reduction in growth at a final pH of 4.69 (95% CI: 4.52-4.96, **Figure 6C**).

**Figure 6.**
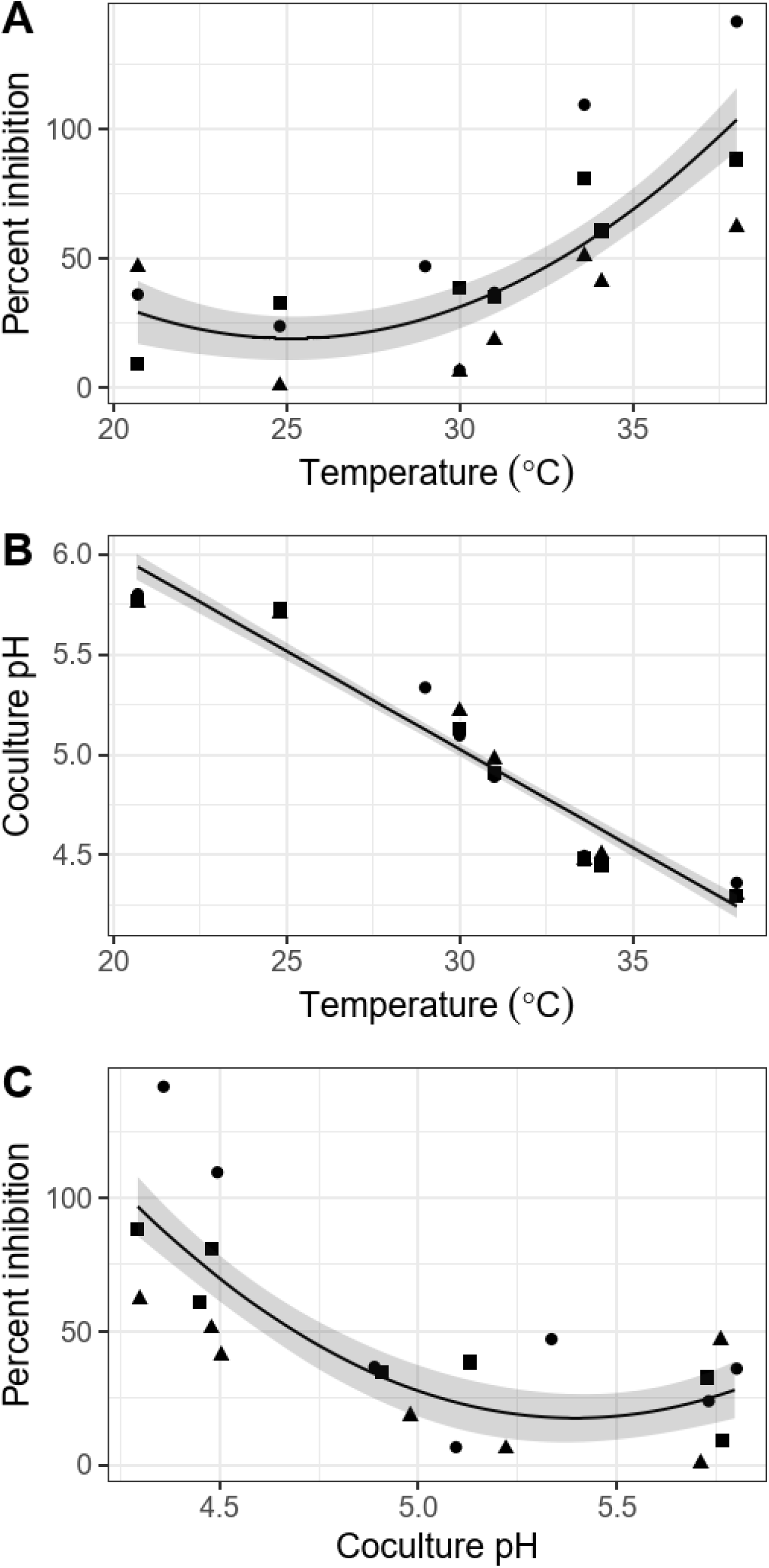
Temperature-dependent, pH-associated effects of *Lactobacillus* on growth of *Lotmaria passim* in symbiont-parasite cocultures. **(A)** Proportional inhibitory effects of *Lactobacillus* on *L. passim* increased with temperature. **(B)** Higher temperatures resulted in lower pH of the growth media. **(C)** Lower pH was correlated with greater proportional inhibition of *L. passim* growth. Lines and shaded bands show estimated marginal means and standard errors of linear model predictions. Points show individual observations. Shapes indicate different experimental blocks.

## DISCUSSION

Due to the fundamental effects of temperature on biological rates, including parasite and symbiont growth and metabolism, temperature has the potential to alter the effects of symbionts on parasite establishment, such that both symbiont and parasite physiology shape the temperature dependence of infection. We found that high temperatures augmented growth rates of symbionts relative to parasites, that symbionts inhibited parasite growth via production of acids, and that high temperatures amplified this inhibitory effect.

### Symbionts were more heat-tolerant than parasites

Our finding of greater heat tolerance in the honey bee symbiont than in the trypanosomatids is consistent with differences in conventional culturing temperatures and results from bumble and honey bee-associated microbes. Most trypanosomatids are cultured at 25-28 °C (63), whereas honey bee gut bacteria thrive at 35-37 °C (22). The ∼6 °C lower peak growth temperature of *L. passim* relative to the *Lactobacillus* symbionts closely resembles the difference in peak temperatures for *C. bombi* (∼34 °C) and *Lactobacillus bombicola* (∼40 °C) from bumble bees (44). Other key symbionts of honey bees are also heat-tolerant and able to grow above 40 °C (64). In our experiments, the *Lactobacillus* symbiont’s greater heat tolerance and temperature responsiveness led to rapid changes in the relative growth rates of symbionts vs. parasites, especially the less heat-tolerant *L. passim*, over the temperature range found in colony-dwelling honey bees (20-37 °C (21)). This suggests that honey bees’ endothermic behavior could select for growth of coevolved symbionts at the expense of opportunistic parasites. Although both trypanosomatids were still able to grow above 34 °C, growth rates of *Lactobacillus* relative to the parasites (especially *L. passim*) increased particularly sharply over the 33.8-37 °C range of the colony’s core during the brood-rearing season (20). This indicates that the warm temperatures at the center of the colony would favor formation of a core symbiont-dominated microbiome during the microbiome’s formative period in the 2-4 days after emergence of adults from pupation, including primacy of *Lactobacillus* Firm-5 species in the rectum (41).

### Symbiont-produced acids inhibited parasite growth

Although growth rates depicted suitability of the colony habitat for trypanosomatids relative to symbionts, growth of symbionts is only directly relevant to proliferation of parasites if there is some interaction between the two. We found that *Lactobacillus-*spent medium inhibited growth of both honey bee parasites in an acidity-dependent manner within the gut pH range observed in bees. Environmental pH is a strong driver of gut microbiome composition (65), where single-unit changes in pH alter the relative growth rates of cooccurring species and community-level production of short-chain fatty acids (66). Production of acids by fermentative bacteria, including Lactobacilli, contributes strongly to their chemical inhibition of gut parasites (67, 68). In humans, diets that acidify stool pH from 6.5 to 5.5 are associated with suppression of opportunistically pathogenic Enterobacteriaceae species, such as *E. coli*, that grow poorly in acidic environments (69). Similarly, supplementation of infants with milk sugar-fermenting *Bifidobacteria* reduces stool pH from 5.97 to 5.15 and reduces levels of Enterobacteriaceae, Clostridiaceae, and endotoxins (70), whereas reduced levels of Bifidobacteria have been associated with a 1.5-unit increase in fecal pH and increased abundance of Enterobacteriaceae, Clostridiaceae, and other dysbiosis-associated bacteria over the past century (71).

In the gut of honey bees, one effect of bacterial colonization is a nearly identical reduction in hindgut pH, from pH ∼6 to pH ∼5.2 (57). A major contributor to this acidification is likely the *Lactobacillus* Firm-5 species cluster, which alone was responsible for around 90% of the chemical changes seen in the gut metabolome of honey bees with intact microbial communities (43). Our findings of *Lactobacillus* acidity-mediated inhibition of honey bee trypanosomatids are consistent with pH-dependent inhibition of *C. bombi* by *Lactobacillus bombicola* from bumble bees (45), and match the negative correlations between *C. bombi* infection intensity and abundances of acid-producing *Lactobacillus* and *Gilliamella* in these hosts (38, 40). Our results also show a strong correlation between pH and *L. passim* growth in cocultures, with inhibition occurring over a physiologically relevant range.

The rectal pH of adult bees in colonies ranged between 4.2 and 6.0 (72). Whereas pre-colonization of bees with *Snodgrassella alvi*—which consumes organic acids (43)—increased *L. passim* infection (48), we found that an endpoint coculture pH below 5 was associated with sharply reduced *L. passim* growth rates, suggesting that microbial or climatic factors that acidify gut pH to the lower end of this observed range would improve resistance to infection. Such acidity-driven pressure on parasites could be one explanation for honey bee parasites’ high tolerance of acidity relative to trypanosomatids from insects with less acidic guts (51).

### High temperatures amplified the parasite-inhibiting effects of symbionts

Our coculture experiments showed that the inhibitory effects of symbionts on parasites increased with temperature. Both *Lactobacillus* strains grew faster and produced acids more rapidly as temperatures increased over the range found in bee colonies. This high-temperature inhibition resulted in lower peak parasite growth rates and reduced parasite heat tolerance relative to when symbionts were absent. Hence, social bees’ endothermic maintenance of high colony temperatures could exert antiparasitic effects not only through direct parasite inhibition, but also via potentiation of the parasite-inhibiting effects of symbionts. The bumble bee symbiont *Lactobacillus bombicola* resulted in similar temperature-dependent inhibition of the parasite *C. bombi,* lowering the temperature of peak parasite growth rate and resulting in a more rapid decline in parasite growth rate as temperatures increased (44). Using a parasite-symbiont system from a related host, our results build on these findings by investigating a broader range and increased number of temperatures, enabling formal comparisons between parasite thermal performance curves in the presence vs. absence of symbionts.

The manipulability of both microbiome and body temperature in a host with a predominantly endothermic life history strategy makes honey bees a useful system to explore the value of endothermy as a reinforcer of symbioses and symbiont-mediated defense against parasites. Prior work has shown that high (35 °C) temperatures increase resistance of honey bees to colonization by heat-sensitive environmental yeasts relative to cooler (29 °C) temperatures, consistent with the seasonal appearance of yeasts during colder weather in field-collected forager bees (73). On the other hand, high (35 °C) temperatures enhanced growth of and gut colonization by the core symbiont *S. alvi* in bumble bees relative to low (28-29 °) temperatures (64). These results implicate bee endothermy in both resistance to opportunistic gut microbes and establishment of key symbionts.

Our results indicate that higher temperatures not only promote symbiont growth, but also augment symbionts’ value as a defense against gut parasites. Consistent with this idea, honey bees collected from cooler winter colonies exhibit gut colonization by non-core microbial taxa in addition to— rather than in place of—their pre-existing core symbionts (42), suggesting that the core symbiont community’s resistance to invasion requires high temperatures even when its major members, including *Lactobacillus* Firm-5, are abundant (42). Temperature-mediated inhibition of trypanosomatids specifically is of practical importance for honey bees, where infection intensities were highest mid-winter in one longitudinal survey (36) and correlated with winter colony collapse in another (26). Environmental factors that perturb thermoregulation, such as exposure to pesticides, agricultural landscapes, and food shortage (21, 74–76), could exacerbate this seasonal susceptibility.

## CONCLUSIONS

The presence of parasites can shift the thermal niche of hosts towards temperatures that enhance escape from or suppression of infection (15). High body temperatures often mitigate infection of warm-adapted hosts, including honey bees and other insects (10, 77–80), with less heat-adapted parasites (8). Conversely, the host’s responses to temperature can alter the optimal temperature for parasite reproduction (5, 8). Our findings complement these results, showing how the presence of gut symbionts can limit both peak growth rate and heat tolerance of a globally prevalent honey bee gut parasite. The extent of symbiont-mediated inhibition varied substantially over the range of temperatures and gut pH levels previously reported in adult bees, indicating the potential for endothermic behavior and core gut symbionts of honey bees to synergistically inhibit parasite growth. Our finding that high temperatures promote symbiont-mediated parasite inhibition shows how changes in host temperature could augment or diminish the protective effects of the microbiome against infection. This illustrates an interaction between the well-established effects of temperature and gut microbial communities on resistance to parasites, with implications for understanding the effects of climate on infection of ectotherms and the evolution of endothermy and fever. The ability of symbionts to shift the thermal niche of parasites highlights the value of investigating parasite thermotolerance in the context of the host microbiota.

## ACKNOWLEDGMENTS

We thank the ATCC, R.S. Schwarz, J.E. Powell, and N.J. Moran for microbial strains and culturing advice; Daniel Padfield for R scripts; and anonymous reviewers for their service in improving the manuscript.

## FUNDING

This project was supported by USDA-NIFA Postdoctoral Fellowship 2022-67012-37482 to ECPY; USDA-NIFA Pollinator Health Grant 2020-67013-31861 to JDE and ECPY; a North American Pollinator Protection Campaign Honey Bee Health Improvement Project Grant and an Eva Crane Trust Grant to ECPY and JDE, and the USDA Agricultural Research Service Beltsville Bee Research Laboratory in house fund. Funders had no role in study design, data collection and interpretation, or publication. We thank the reviewers for their service in improving the manuscript.

## CONFLICTS OF INTEREST

The authors declare that they have no conflicts of interest.

## DATA AVAILABILITY

All data are supplied in the Supplementary Information, Data S1.

## AUTHORS’ CONTRIBUTIONS

ECPY conceived the study. ECPY and LM designed experiments. ECPY, LM, and WFH conducted experiments. ECPY analyzed data. ECPY and LM drafted the manuscript with guidance from JDE. All authors revised the manuscript and gave approval for publication.

## Supplementary information

### SUPPLEMENTARY FIGURES

**Supplementary Figure 1.**
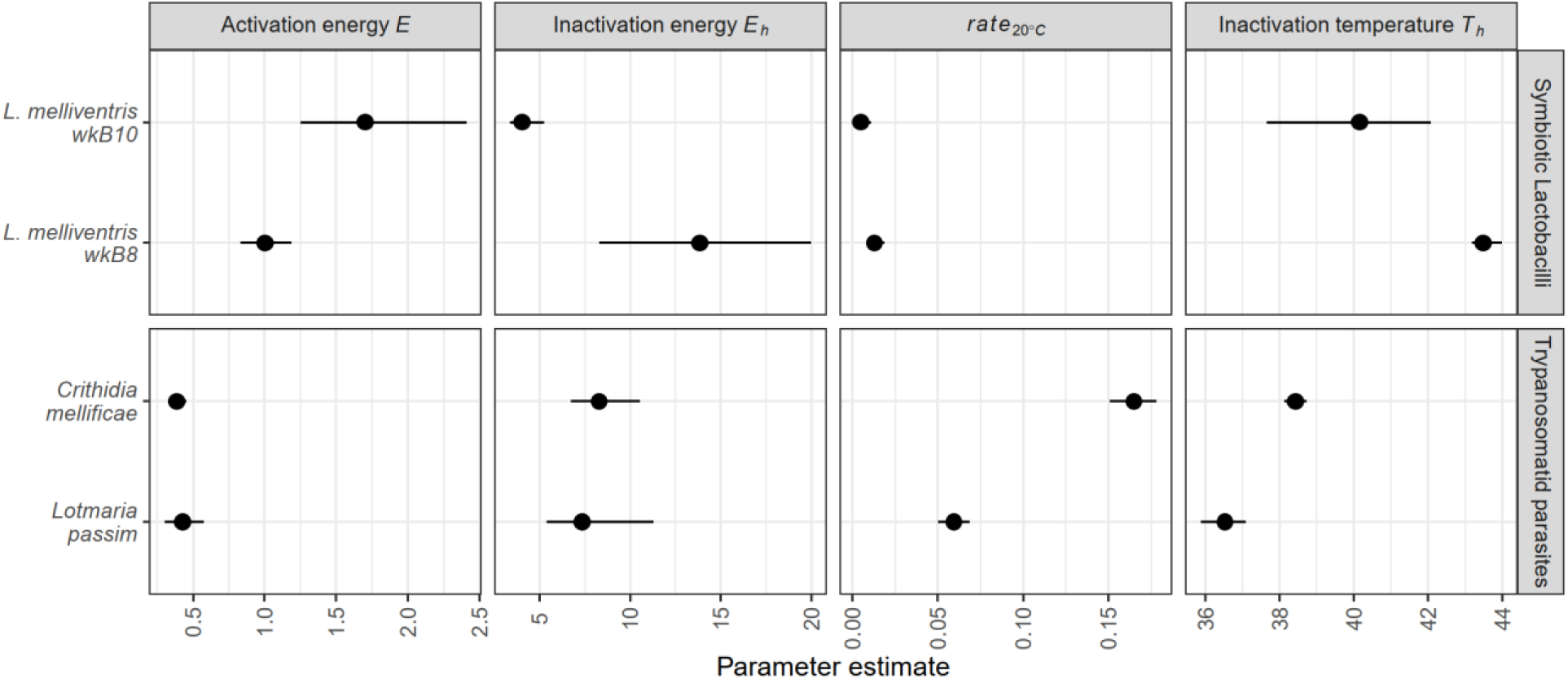
Model parameter estimates for Sharpe-Schoolfield models. Parameters *E* and *E_h_* (measured in eV) indicate how strongly growth rate responds to temperature below and above the temperature of peak growth, respectively. The growth rate at 20 °C (measured in h^-1^) indicates the speed of growth at a reference temperature at which no high-temperature inhibition of growth is expected. Parameter *T_h_* (calculated in K, shown in °C) indicates the temperature at which growth rate is inhibited by 50% relative to the rate predicted by the upward-sloping (Arrhenius) portion of the thermal performance curve, which assumes a monotonic increase of growth rate with increasing temperature. Points and error bars show bootstrap medians and 95% confidence intervals.

**Supplementary Figure 2.**
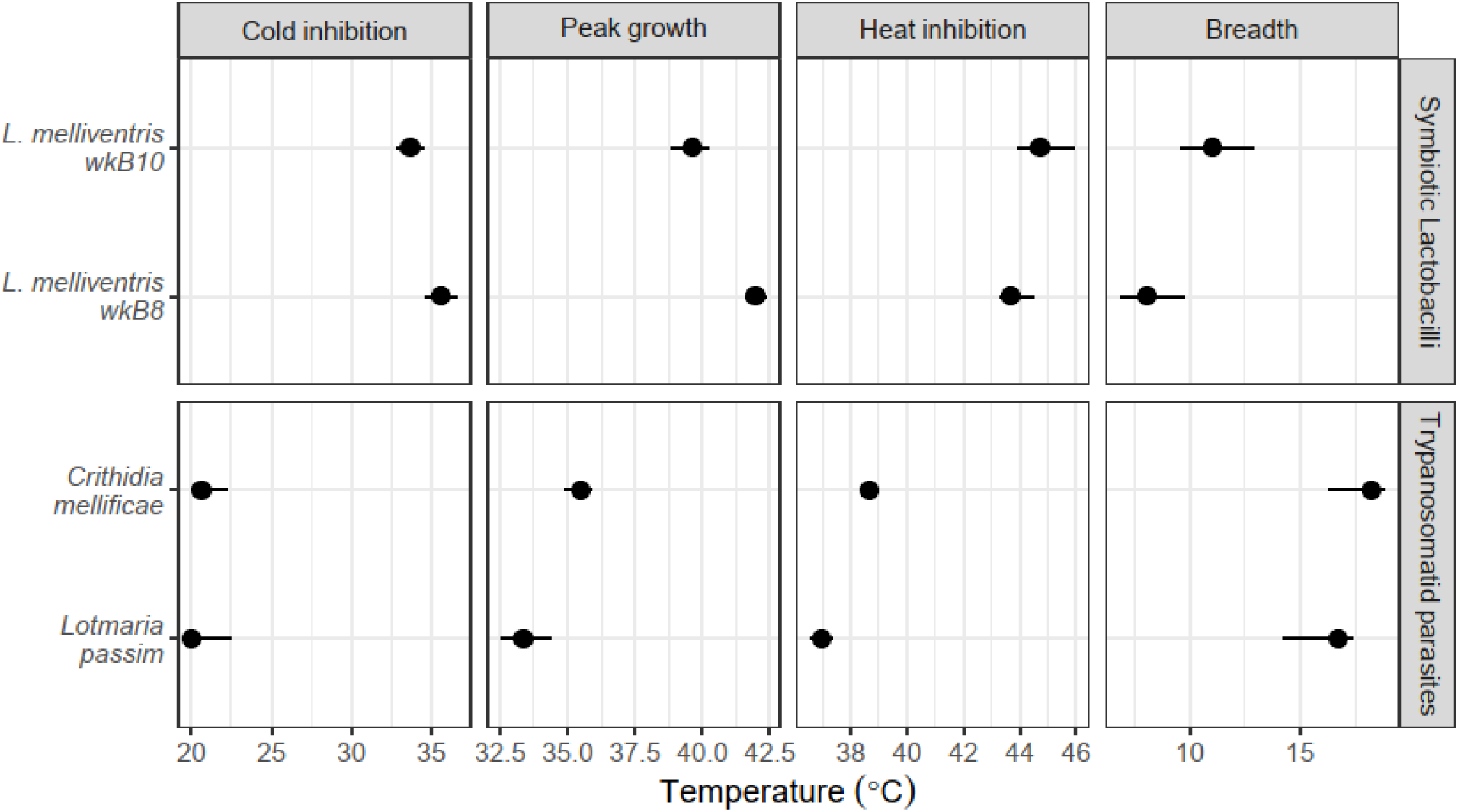
Estimates of derived quantities from Sharpe-Schoolfield models for temperatures of peak growth, 50% inhibition due to cold and hot temperatures, and thermal niche breadth. Panel for “Peak growth” shows the temperature at which peak growth rate occurs. Panels for “Cold inhibition” and “Heat inhibition” show the temperatures at which growth is reduced by 50% relative to the peak growth rate. “Breadth” indicates the size of the temperature interval over which growth rate is at least half of maximum, calculated as the difference between the temperatures of cold and heat inhibition. Points and error bars show bootstrap medians and 95% confidence intervals.

**Supplementary Figure 3.**
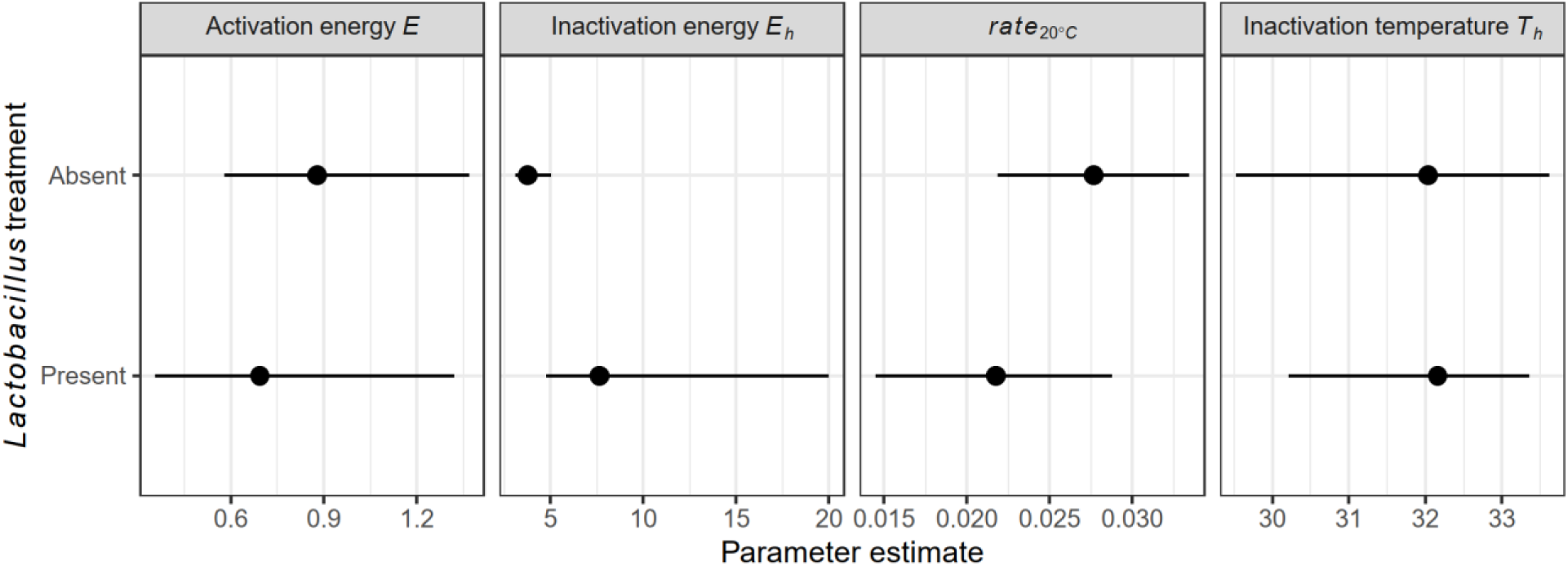
Model parameter estimates for Sharpe-Schoolfield models for *Lotmaria passim* growth in the presence and absence of *Lactobacillus*. Parameters *E* and *E_h_* (measured in eV) indicate how strongly growth rate responds to temperature below and above the temperature of peak growth, respectively. The growth rate at 20 °C (measured in h^-1^) indicates the speed of growth at a reference temperature at which no high-temperature inhibition of growth is expected. Parameter *T_h_* (calculated in K, shown in °C) indicates the temperature at which growth rate is inhibited by 50% relative to the rate predicted by the upward-sloping (Arrhenius) portion of the thermal performance curve, which assumes a monotonic increase of growth rate with increasing temperature. Points and error bars show bootstrap medians and 95% confidence intervals.

**Supplementary Figure 4.**
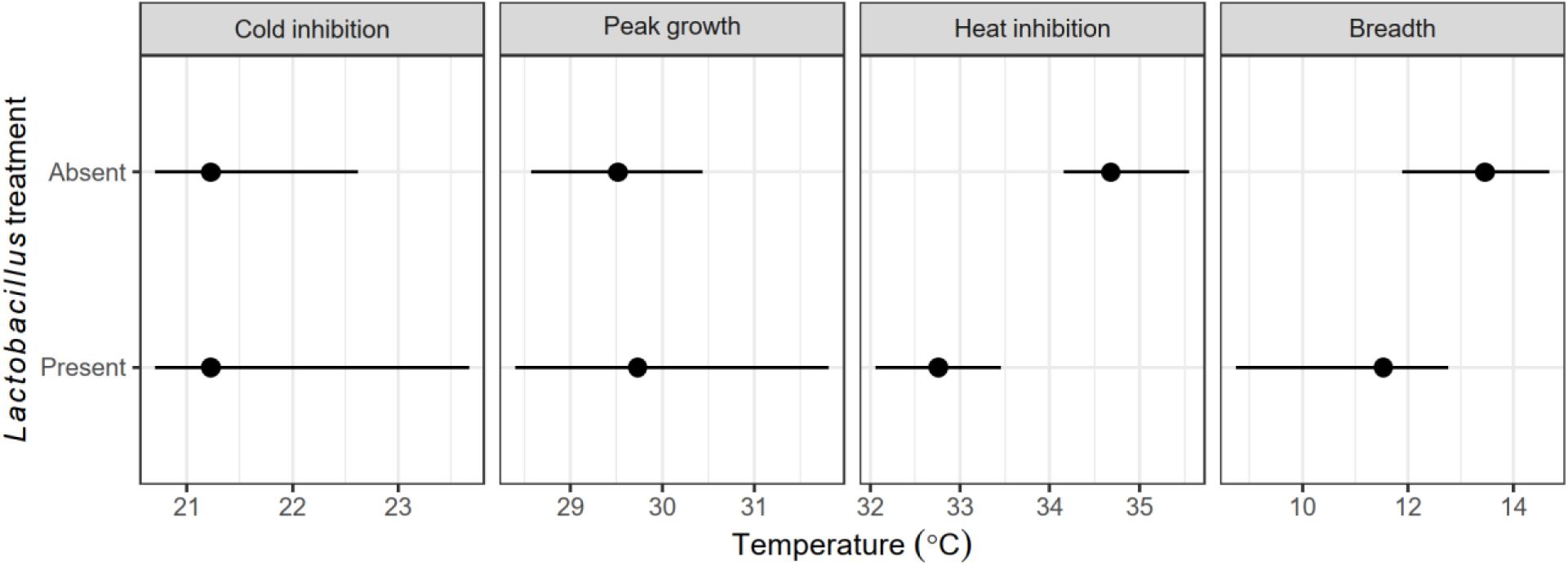
Estimates of derived quantities from Sharpe-Schoolfield models of *L. passim* growth in the presence and absence of *Lactobacillus.* Panels for “Cold inhibition” and “Heat inhibition” show the temperatures at which growth is reduced by 50% relative to the peak growth rate. “Breadth” indicates the size of the temperature interval over which growth rate is at least half of maximum, calculated as the difference between the temperatures of cold and heat inhibition. Points and error bars show bootstrap medians and 95% confidence intervals.

### SUPPLEMENTARY DATA

**Supplementary Data S1 (.zip). Zipped file containing raw data** with separate text files for growth rates of *Lactobacillus* and trypanosomatids in isolation; growth of trypanosomatids with *Lactobacillus* spent medium; growth of *L. passim* in cocultures; and summaries of Sharpe-Schoolfield models for *Lactobacillus* and trypanosomatids in isolation and *L. passim -Lactobacillus* cocultures.

## Notes

### Competing Interest Statement

The authors have declared no competing interest.

